# Heteroscedastic Personalized Regression Unveils Genetic Basis of Alzheimer’s Disease Stratified by Cognitive Level

**DOI:** 10.1101/2023.09.12.557499

**Authors:** Zhirong Chen, Haohan Wang

## Abstract

In contemporary medical research, patient heterogeneity plays a pivotal role in comprehending intricate diseases such as Alzheimer’s disease and various forms of cancer. Specifically, in the genomic analysis of Alzheimer’s disease, individual patients may exhibit unique causal mutations that significantly influence their therapeutic trajectory. Conventional models that share numerous parameters across all individuals struggle to discern this heterogeneity and identify the influential factors for individuals. To tackle this challenge, we propose an innovative approach called Heteroscedastic Personalized Regression (Het-PR) to estimate the heterogeneity across samples and obtain personalized models for each sample. We demonstrate the effectiveness and robustness of Het-PR through both simulation and real data experiments. In the simulation experiment, we show that Het-PR outperforms other state-of-the-art models in capturing inter-sample heterogeneity. In the real data experiment, we apply Het-PR to Alzheimer’s data and show that it can identify persuasive selected genetic factors for each individual patient. Interestingly, our results suggest that there might be different associative SNPs for AD patients stratified by different cognitive levels.

**Author summary:** In medical research, it has been observed that causes of a disease vary significantly among individuals, especially when looking at complex diseases like Alzheimer’s disease and cancer. For Alzheimer’s disease, obesity, age, gender, and depression may play different roles across different patients. When studying the genes of Alzheimer’s patients, we find that each person might have their own unique genetic changes that can affect their treatment. For example, Alzheimer’s patients with different genetic mutations may respond differently to the same treatment. Traditional research methods often miss these individual differences and can’t always pinpoint important personalized factors for each patient, because they usually use one model for all patients. To better understand these differences, we’ve introduced a new method, Heteroscedastic Personalized Regression (Het-PR), which generates a personalized model for each individual. Our experiments show that Het-PR is more effective than other leading methods in identifying these patient differences and recognizing Alzheimer’s genetic basis for each patient through both simulation and real data experiments. When we used Het-PR on real Alzheimer’s data, it helped us spot key genetic factors for each patient. Additionally, in our study, we excitedly find that different genetic markers in Alzheimer’s patients are possibly based on their cognitive abilities. Software for Heteroscedastic Personalized Regression is available in https://github.com/rong-hash/Het-PR.

## Introduction

The prevalence of heterogeneous diseases, such as Alzheimer’s and cancer, is increasing across patients [1, 2]. For example, Alzheimer’s disease is influenced by various factors including gender, mid-life obesity, depression, and others, which contribute to significant disparities in patients’ clinical manifestations [3, 4]. Among all latent factors, genetic basis, such as SNP, is one of the important heterogeneous factors that influence disease progress, treatment response, and clinical trial design. For instance, APOE *ϵ*4 carriers may respond differently to certain treatments compared to non-carriers [5]. What’s more, as for the influence on disease progress, a study in 2021 showed that APOE *ϵ*4 carriers and individuals with abnormal baseline tau levels showed a faster decline at the group level, but also greater within-group variability [6]. Consequently, understanding this heterogeneity is imperative for the development of personalized treatment strategies and the design of efficacious clinical trials. To comprehensively understand this heterogeneity, it is essential to construct a personalized profile for each patient.

While conventional models estimate a single set of common parameters for all samples, personalized regression models offer several advantages in uncovering heterogeneity: 1) These models can identify specific and informative factors relevant to individual patients, thus providing more refined insights compared to traditional machine learning models. 2) Upon identifying the key contributing factors for individual patients, it becomes feasible to develop tailored therapeutic strategies that address each patient’s unique characteristics. 3) Personalized regression models only take a small amount of data to build a personal model for each patient.

With the incorporation of personalized effects, the model becomes more interpretable and can offer valuable insights into clinical treatment and pathological analysis. In fact, personalized models have also been shown to solve problems in various domains, including finance, society, and more [7, 8].

Prior studies have illustrated the potential of diverse personalized regression models in the medical field. Conventional personalized regression models predominantly employ penalized terms to achieve personalization for individual samples. For example, Localized Lasso [9] extends the regularization term of Network Lasso [10] to facilitate personalized effects; Penalized Angular Regression accomplishes personalization by establishing distinct penalized terms for each sample [11]; Personalized Regression with Distance Matching achieves personalization by matching covariate distances and personalized parameter distances [7, 8]. While previous personalized models have demonstrated effectiveness in prediction tasks, their performance in feature selection tasks remains suboptimal, thus lacking sufficient support for the physiological analysis of patients.

Moreover, a variety of techniques generally employ a similar set of principles to regularize the coefficients of related samples. Some of these methods were developed prior to personalized regression models. Examples include varying-coefficient models [12–17] and contextualized models [18–20].

However, these methods typically face two primary challenges: 1. The specifically designed regularization to express the relationship restricts the model’s applicability to other scenarios; 2. The estimation of the model parameters for a single sample usually relies primarily on a subset of samples, which limits the statistical efficiency of the estimation process. 3. The models usually need a large amount of data to achieve personalized effect, which is hard to access in the real world.

To address the above issues, we propose a novel personalized regression framework that employs heteroscedasticity (Het) assumption through a Linear Mixed Model (LMM) construction for estimating individualized parameters of each patient. This approach diverges from the conventional personalization methods that rely on regularization. In our proposed framework, we construct an LMM for each sample based on the distance among the sample features. To be more specific, given a matrix of data with dimensions *n*× *p*, where *n* denotes the number of samples and *p* represents the number of features, Personalized Regression can be trained with this data to estimate a set of parameters of dimensions *n* × *p*, for example, *p*-value of the importance of each feature associated with the label for feature section. By sorting the *p*-values and establishing a threshold, the top-ranked features can be extracted for further study with potential relationships with the labels.

To assess the efficacy of the proposed method, we carried out both simulation and real data tests. Our simulation tests indicated that our model exhibited exceptional performance in the AUCROC score, showcasing its robust variable selection capabilities. In the real data test, we applied our model to the Alzheimer’s Disease Neuroimaging Initiative (ADNI) database. We found that the Single Nucleotide Polymorphisms (SNPs) extracted for each patient displayed high interpretability and reliability. Notably, many genes corresponding to the selected SNPs had been previously validated as Alzheimer’s-associated genes. Furthermore, the extracted SNPs exhibited a strong correlation with the patients’ cognitive levels, underlining the potential of our model in facilitating the development of personalized medical treatment plans.

## Methods

### Background -Linear Regression

To elucidate the methodology underlying the Het-PR model, we begin with the notation and the simple linear regression model.

Given the dataset of (**X, Y**) where **X** ∈ ℝ^*n×p*^ and **Y** ∈ ℝ^*n*^, where *p < n*, and (**x, y**) to denote one sample of (**X, Y**), we use subscripts to denote samples and superscripts to denote features, thus 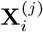 denotes the *j*^*th*^ feature of the *i*^*th*^ sample.

Linear regression assumes the data generation process of

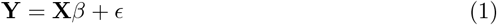

where *β* is the *p* dimension coefficients, and *ϵ* denotes a vector of length *n* of residue error.

It is well known in the community that, if *ϵ* ∽ *N* (0, *σ*^2^), we can have

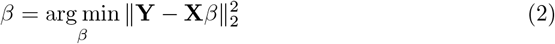

as an unbiased estimator.

### Background -Personalized Regression

For Personalized Regression, instead of sharing the parameter *β* among all the samples, we assign a particular parameter set for each sample, which can be shown as

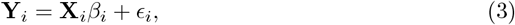

where *i* represents the *i*^th^ sample.

The major differences between personalized regression and linear regression are that we now assume there is a coefficient vector associated with each sample, for which we use *β*_*i*_ to denote the coefficients associated with the *i*^th^ sample. Further, if we use a matrix **B** ∈ ℝ^*n×p*^ to denote the coefficients for all samples, we can have 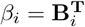.

However, it will be almost impossible to use the traditional MLE-based method i.e., (2)to solve the personalized regression problem (3) because now we only have one sample for every *p* coefficients. Fortunately, previous studies have demonstrated several solutions to estimate the coefficients in such cases with additional assumptions. One assumption that Lengerich [7] uses is that closely related samples will have similar coefficients. Therefore, the problem is formulated as follows.

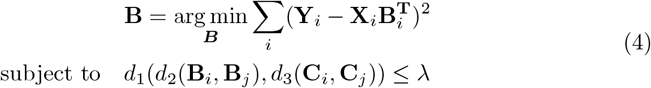

where *d*_1_, *d*_2_, and *d*_3_ are three customized distance functions and **C**_*i*_, **C**_*j*_ are covariates such as demographic information. *λ* is another hyperparameter. With this setup, similar samples will have similar covariates because of the constraint.

As an example, we can consider a more specific form of the above formulation (4) with *d*_1_, *d*_2_, and *d*_3_ assigned concrete choices:

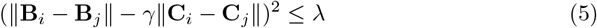

With this specific form, if we can write the main neighborhood assumption in [7] back to the data generation process, (4) estimates the parameters of the following data generation process:

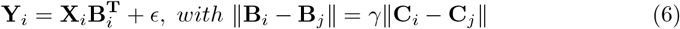

where we again assume 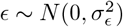, meaning the above problem is a homoscedasticity problem.

In a loosely speaking manner, we can consider the problem in (4) as a “heterogeneous” problem because of the variation introduced through the coefficients for different individuals, but it is homoscedastic because the error term associated with each individual is of the same variance.

### Heteroscedastic Personalized Regression

In this section, we start to introduce our new method. Loosely speaking, our core idea lies in an exchange of variations introduced in linear regression. In contrast to (4), which is a heterogeneous and homoscedastic problem, we formalize the personalized regression as a homogeneous and heteroscedastic one. This new formalization offers us a more efficient manner in estimating the coefficients for each individual, and also comes with extended flexibility to immediately turn different MLE-based regression methods into their personalized version.

### Design Rationale

We first introduce the design rationale of our method by mismatching the sample of interest to the coefficient interest. For example, we study (**X**_*j*_, **Y**_*j*_), but with 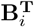, so, we have

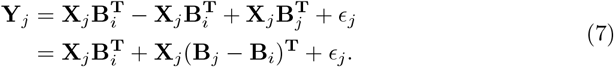

Following our heteroscedasticity view of the problem and assuming the differences introduced from **B**_*j*_ − **B**_*i*_ can be modelled as a noise term, we can continue to substitute (**B**_*j*_ − **B**_*i*_)^**T**^ with **u**_*j*_, and we can get

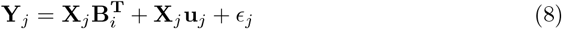

where **u**_*j*_ satisfies 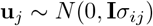, and *σ*_*ij*_ is *d*(**B**_*i*_, **B**_*j*_), where *d* is the customized distance function. For simplicity, here we use the same concrete form discussed in the background section, therefore we have

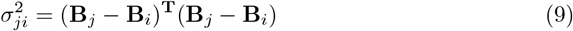

If we continue with the data generation process in (6), we can immediately get 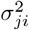 as

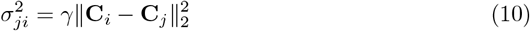

The above derivation shows that we can use all the samples to estimate the coefficients **B**_*i*_ even though **Y**_*j*_ are generated with **B**_*j*_. The main disparity introduced by such a mismatching is the differences raised from **X**_*j*_**B**_*j*_ and **X**_*j*_**B**_*i*_, and we consider this disparity as an error term associated with sample *j*. In other words, all the samples can be used to estimate the coefficients of sample *j*, if the error term for every individual is constructed differently. Thus, loosely speaking, we turn the personalized regression problem into a “homogeneous” but heteroscedastic problem.

### Heteroscedastic Personalized Linear Regression Model

With the above setup, for sample *j*, we have

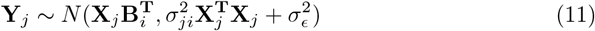

where we do not require any explicit relationship between *j* and *i*. Therefore, to estimate the coefficient of any samples (e.g., sample *i*) we can use all the samples together. Thus, we have:

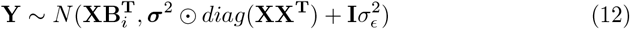

where ⊙ is element-wise product, and ***σ***^2^ is a vector of length *n* with its *j*^*th*^ element to be 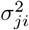.

Conveniently, following the data generation assumption, although 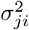 has *n* elements, it only has one parameter *γ*, the rest are determinable through the knowledge of **C**.

As one might notice, (12) is a standard form of linear mixed model (LMM). This convenient connection back to the linear mixed model will allow us immediately to re-use all the mature linear mixed model estimation algorithms to estimate the coefficients of our model, once the error term is constructed properly for the sample of interest. In this work, we use the FaSTLMM algorithm [21] through the implementation of [22]. It is also worth explicitly mentioning that, for every sample of interest, our algorithm needs to construct the error term and perform the estimation. Thus, it requires *n* runs if there are *n* samples of interest for personalized estimation.

### Heteroscedastic Personalized Model Framework

Another significance of our method is that for every population-based model, denoted as

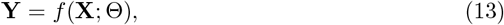

where *f* (·; Θ) is a function with Θ denoting the parameters. We can reuse our above idea to turn it into its personalized version with

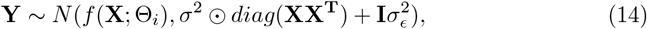

which allows us to estimate the personalized coefficients for sample *i*.

In the experiment section, we will refer to *f* (**X**; Θ) as the “base model” and explore two choices.

## Results

### Simulation Test

#### Simulation Data Generation

We first test our methods in comparison to other methods with simulation data. We start with a simpler situation, where instead of the situations where every individual has one’s own set of oracle effect sizes, we group the individuals into different cohorts, and individuals of the same cohort will have similar parameters. We generate the simulation data with the following process. We first sample *n* samples, each with *p* features from a uniform distribution. We denote the data as **X**. These samples are evenly split into *g* cohorts, and each cohort will have its unique effect size vector *β* with the same sparsity level (*k* features will have non-zero effect sizes). Each of the *k* non-zero values of the *β* vector will be sampled as follows: We first sample a scalar *a* ∈ *N* (0, 1), then

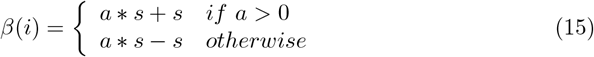

where s is a scaling factor controlling the effect sizes.

Samples within the same cohort will share the same effect size *β*. Thus, for the *i*^*th*^ sample, the label **Y**_*i*_, will be

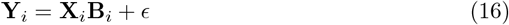

where 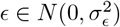 and following the notation of this manuscript, we use **B** to denote the matrix of effect sizes *β*. Further, we use *σ*_*ϵ*_ as another hyperparameter that controls the simulation process.

Finally, to test our model’s behavior, we need to pass the covariates of the demographic information. At the simulation state, we simply use the oracle information of the partition information, i.e., the **B**, as the demographic information **C**.

#### Simulation Result

Since we are interested in recovering the genetic basis of the disease patterns, we evaluate these methods for the ability to recover the non-zero coefficients. We test the methods with AUCROC score of comparing estimated “importance” of coefficients to the Oracle coefficient sparsity pattern. In particular, based on the nature of the methods, the “importance” are *p*-values [23] when available or the absolute values of the estimated coefficients when *p*-values are not available. As we discussed before, our Het-PR model can be equipped with different “base model”, and we tested two versions, the “pl pvalue” denotes the Het-PR with a base model that is a variant of linear mixed model that can report *p*-value even for high-dimension case [22], and “pl beta” denotes the Het-PR with a base model of linear regression. dmr model refers to personalized regression with distance matching [7, 8], par refers to penalized angular regression [11], and pr is population regression. Our results demonstrate that the Het-PR model consistently outperforms other models in accurately recovering the original sparsity pattern of the coefficients across a range of parameter values. Specifically, we conducted five repeated simulations with different random seeds for each parameter set, which varied in terms of sample size (*n*), number of features (*p*), Sparsity Level (*k*), and number of cohorts (*BlockNum*).

After setting the hyperparameters to different values and running the simulation experiment with 16 parameter sets, each parameter set with 5 random seeds, we got the overall performance distribution shown in Figure 1, and detailed model performance is shown in Figure 2. We can find our model performs better than other models in most cases.

**Fig 1.**
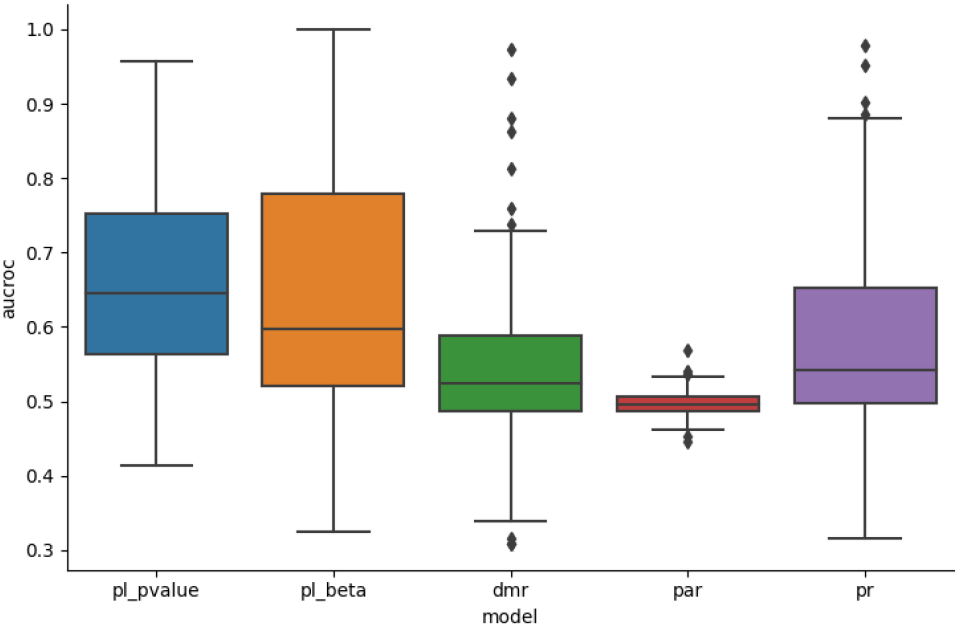
Overall Performance of Simulation Tests across 5 Models. pl pvalue: Personalized Regression with Linear Mixed Model, using p-value to calculate AUCROC score; pl beta: Personalized Regression with Linear Regression, using *β* to calculate AUCROC. dmr: Personalized Regression with Distance Matching; par: Penalized Angular Regression. cr: Clustering Regression; pr: Population Regression

**Fig 2.**
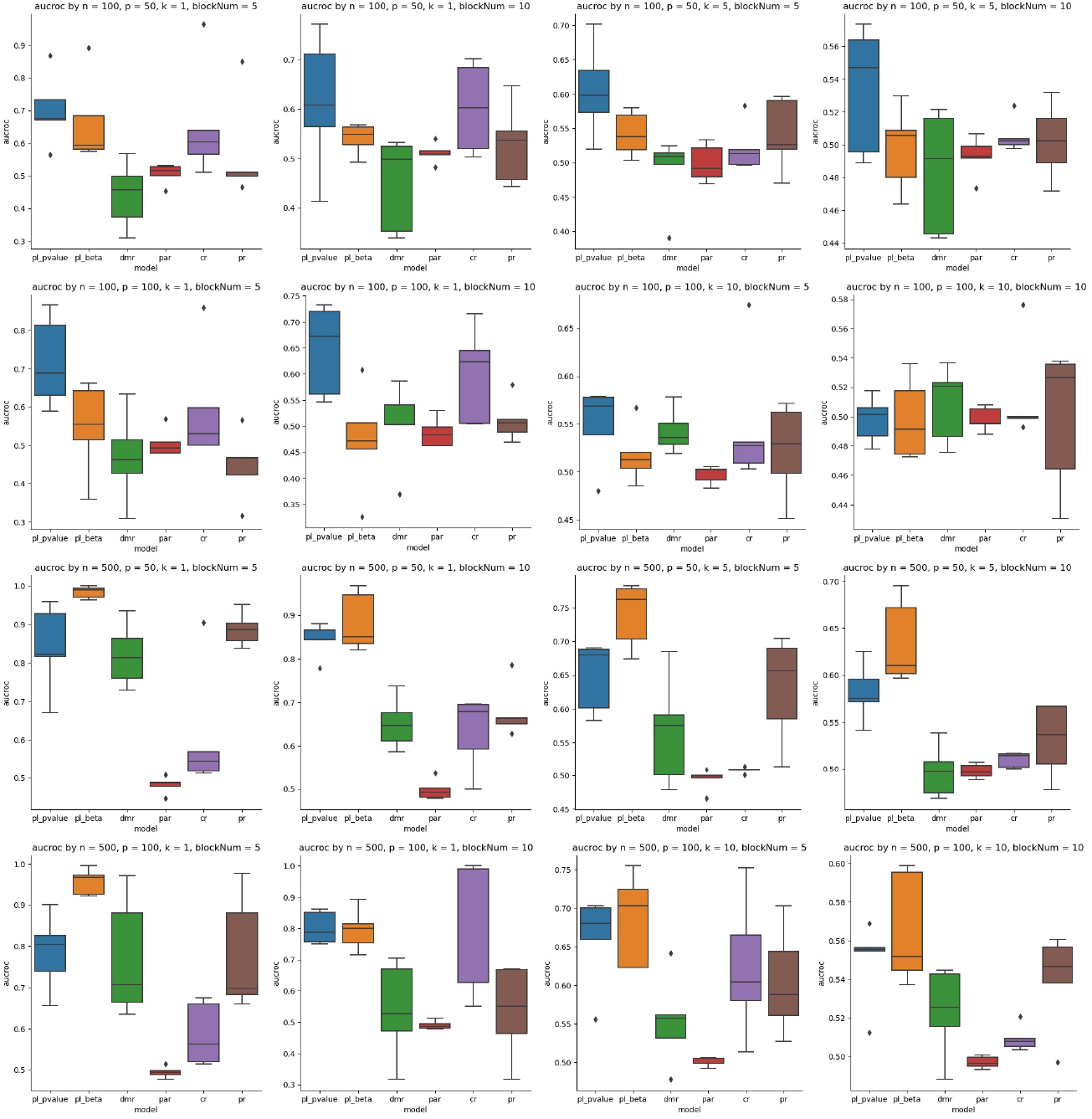
Model Performance under Different Parameters. There are 16 parameter sets in total, each shown in the title of the sub-figure. We conduct five experiments per parameter set with different random seeds.

#### Alzheimer’s Experiment

With the success of simulation experiments, we continue to move on to apply our method to understand the genetic basis of Alzheimer’s disease at a personalized level. Alzheimer’s disease (AD), one of the most lethal ailments affecting the elderly population, has been a focal point of interest for biomedical research communities over several decades. Despite the extensive efforts dedicated to understanding the disease’s mechanisms, effective treatments for AD remain elusive [24–28], owing to the absence of a widely accepted comprehension of its pathology [29]. For instance, it was long theorized that amyloid plaques in the brain were responsible for AD [30–35]. However, this view has been recently contested, suggesting that plaques are merely a result of decreasing levels of soluble amyloid-beta in the brain when normal proteins transform into abnormal amyloid plaques under conditions of biological, metabolic, or infectious stress [36, 37]. Consequently, previous treatments targeting amyloid plaques have become a subject of debate [38].

One factor contributing to the challenge of understanding AD is its heterogeneity. Over the years, researchers have identified several AD subtypes from various perspectives. Genetically, AD is divided into two subtypes: familial AD and sporadic AD [39–43]. However, the genetic basis of sporadic AD is still elusive until now, and a hypothesis behind this challenge is that the genetic pattern of sporadic AD might not be universal across all populations as there are typically three or four subtypes of AD based on tau pathology and brain atrophy [44–47].

From a pathological standpoint, AD is typically classified into three or four subtypes based on tau pathology and brain atrophy [44–47]. Clinically, patterns have emerged, such as the presence of amnestic syndrome, language disturbances, visuospatial skill impairments, attentional function deficiencies, and executive process and praxis disruptions [48]. However, clear associations between subtypes across these different perspectives have not been established [49]. Moreover, AD exhibits a significantly different prevalence between female and male patients, with women affected 1.7 times more frequently than men [4]. This observation implies the existence of additional subtypes when considering gender as a covariate, although previous AD studies have seldom accounted for it.

Therefore, our personalized regression model can potentially identify the genetic patterns of AD by considering its heterogeneity, thus constructing a personalized profile for each patient. We test our model on the ADNI 1 genotype data, which contains 192 patients in total, consisting of 79 patients (sporadic AD) and 113 normal controls. Each sample has 257361 SNPs. Each SNP has 3 alternative values, 0, 1, 2, representing no variation, 1 variation, and 2 variations respectively.

Due to the small sample size, in this study, we only use exon markers as **X** for each sample, and the rest of SNPs as **C**. Thus, we do not have to use covariates explicitly offered by the ADNI in our model but we can save them to evaluate whether the genetic patterns we identified are meaningful. After training on the data using Het-PR, we can get the *p*-value of SNP for each sample, and we consider the threshold of *p*-value as 0.05.

We first report and analyze the results at the population level by collecting the most frequently reported SNPs across all the samples. We report the top 10 SNPs for all patients in Table 1. In Table 1, we find that 7 out of these 10 genes are strongly related to Alzheimer’s [50–56]. Furthermore, although we don’t have strong evidence that the other 3 genes are related to Alzheimer’s, we still find it related to brain activity. For example, for the most selected gene *CFAP73*, it’s an important paralog of the gene *CCDC42*, which is specifically expressed in testis and brain [57]. Overall, the result suggests that our model can identify SNPs implicated with AD.

**Table 1.** Top 10 SNPs across all patients. Alzheimer’s related genes are bolded. NA means this gene is not named yet. Most influential genes are in the protein-coding type.

| SNP | Chromosome | Location | Gene ID | Gene Name | Gene Type |
| --- | --- | --- | --- | --- | --- |
| *rs2701108* | chr12 | 113158644 | ENSG00000186710.12 | *CFAP73* | protein_coding |
| *rs4747905* | chr10 | 11463463 | ENSG00000148429.15 | ***USP6NL*** | protein_coding |
| *rs7856501* | chr9 | 131009377 | ENSG00000050555.19 | ***LAMC3*** | protein_coding |
| *rs1865604* | chr3 | 150741859 | ENSG00000181788.4 | *SIAH2* | protein_coding |
| *rs11633318* | chr15 | 75389753 | ENSG00000169375.17 | ***SIN3A*** | protein_coding |
| *rs12562620* | chr1 | 214353050 | ENSG00000228470.1 | NA | lncRNA |
| *rs4445847* | chr15 | 71959066 | ENSG00000066933.16 | ***MYO9A*** | protein_coding |
| *rs2509172* | chr11 | 69654072 | ENSG00000110092.4 | ***CCND1*** | protein_coding |
| *rs8005728* | chr14 | 20380371 | ENSG00000129566.13 | ***TEP1*** | protein_coding |
| *rs13275051* | chr8 | 70607376 | ENSG00000067167.8 | ***TRAM1*** | protein_coding |

Further, since our model is particularly designed to understand the personalized genetic basis, we further report the results for its ability in unveiling the genetic basis of each patient. For the discussion, we consider the patients whose selected SNPs are more than 10. For each patient, the top 10 genes corresponding to the SNPs are shown in Table 2.

**Table 2.**
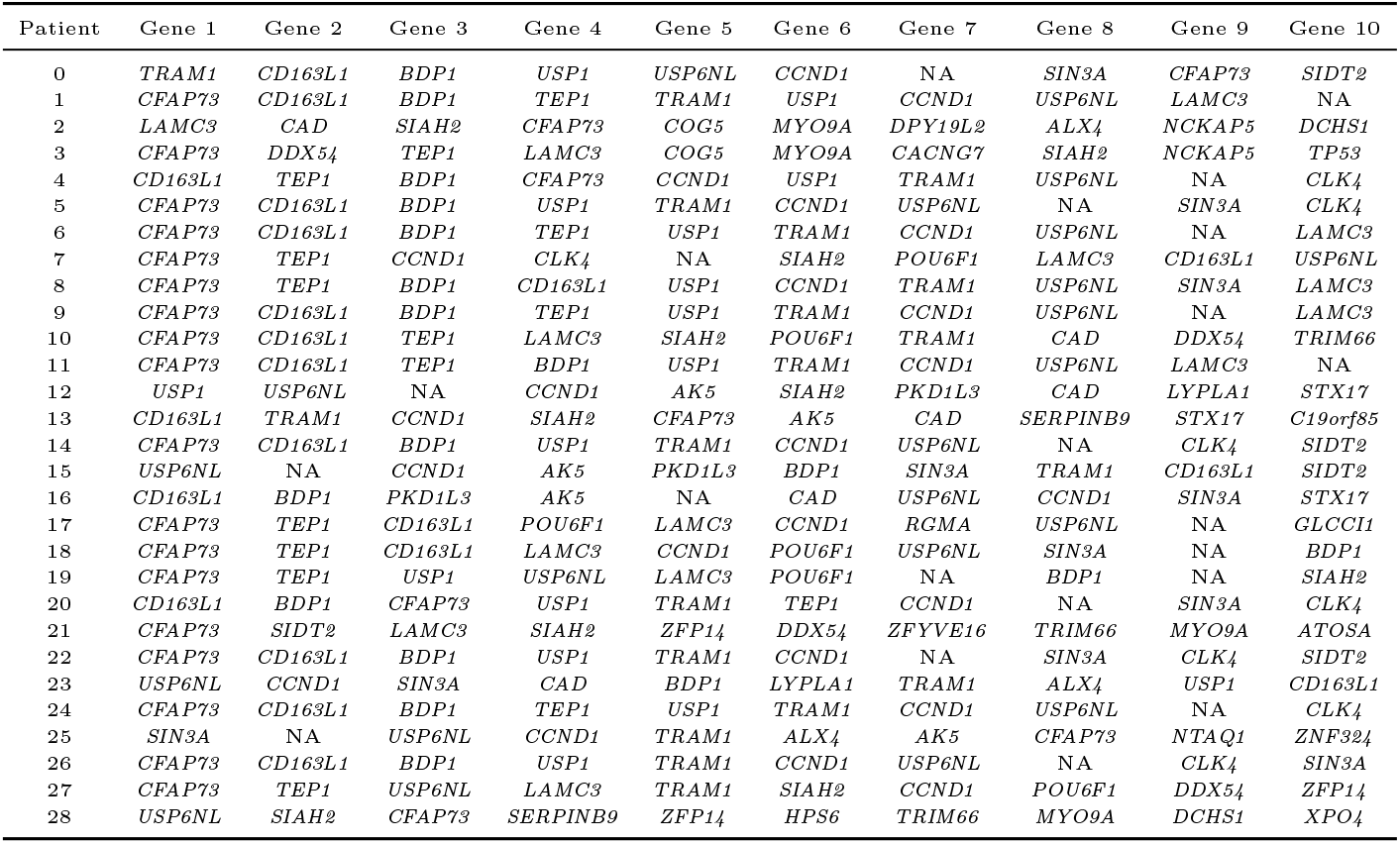
Top 10 influential genes for each patient.

To offer a further understanding of these results, we conduct a hieratical clustering based on the *p*-value of the SNPs, shown as Figure 3. Based on the clustering, we can divide the patients into two groups. Group1 is colored blue and Group2 is colored orange in Figure 3.

**Fig 3.**
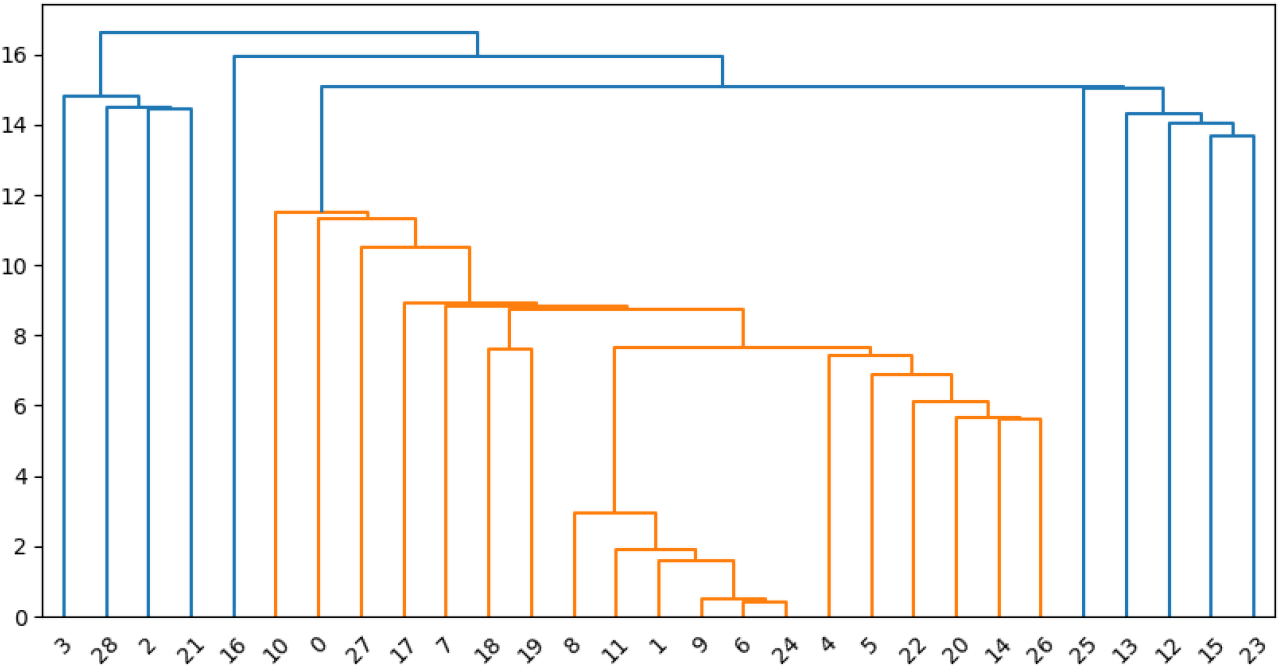
Hieratical Clustering based on pvalue of SNPs, Group1 is colored blue, and Group2 is colored by orange, which is the same as Figure 4.

Interestingly, we notice that this group partition is correlated with the group created by patients’ cognitive level, which can be represented by 8 authoritative targets: CDRSB, ADAS11, ADAS13, ADASQ4, MMSE, RAVLT immediate, RAVLT learning, RAVLT perc forgetting. The meaning of these 8 targets is shown in Table 3

**Table 3.** Cognitive Level Target. 8 cognitive test scores given by ADNI.

| Target | Description | Score |
| --- | --- | --- |
| CDRSB | Clinical Dementia Rating Scale Sum of Boxes | Higher, Severe |
| ADAS11 | Alzheimer's Disease Assessment Scale-Cognitive Subscale 11 | Higher, Severe |
| ADAS13 | Alzheimer's Disease Assessment Scale-Cognitive Subscale 13 | Higher, Severe |
| ADASQ4 | Alzheimer's Disease Assessment Scale-Cognitive Subscale Question 4 | Higher, Severe |
| MMSE | Mini-Mental State Examination | Lower, Severe |
| RAVLT_immediate | immediate recall component of the Rey Auditory Verbal Learning Test | Lower, Severe |
| RAVLT_learning | Rey Auditory Verbal Learning Test | Lower, Severe |
| RAVLT_perc_forgetting | amount of information that is forgotten | Higher, Severe |

We compare the scores between Group1 and Group2 and the result is shown in Figure 4. From the experiment, we find that eight out of eight targets show that the cognitive level of Group1 is lower than Group2, meaning that the selected SNP is strongly related to the patients’ cognitive level. The *p*-value of the t-test is shown in Figure 4. We consider these results very interesting as these discussions might be one of the first results to show that the genetic basis of AD patients might be stratified by the cognitive level of patients.

**Fig 4.**
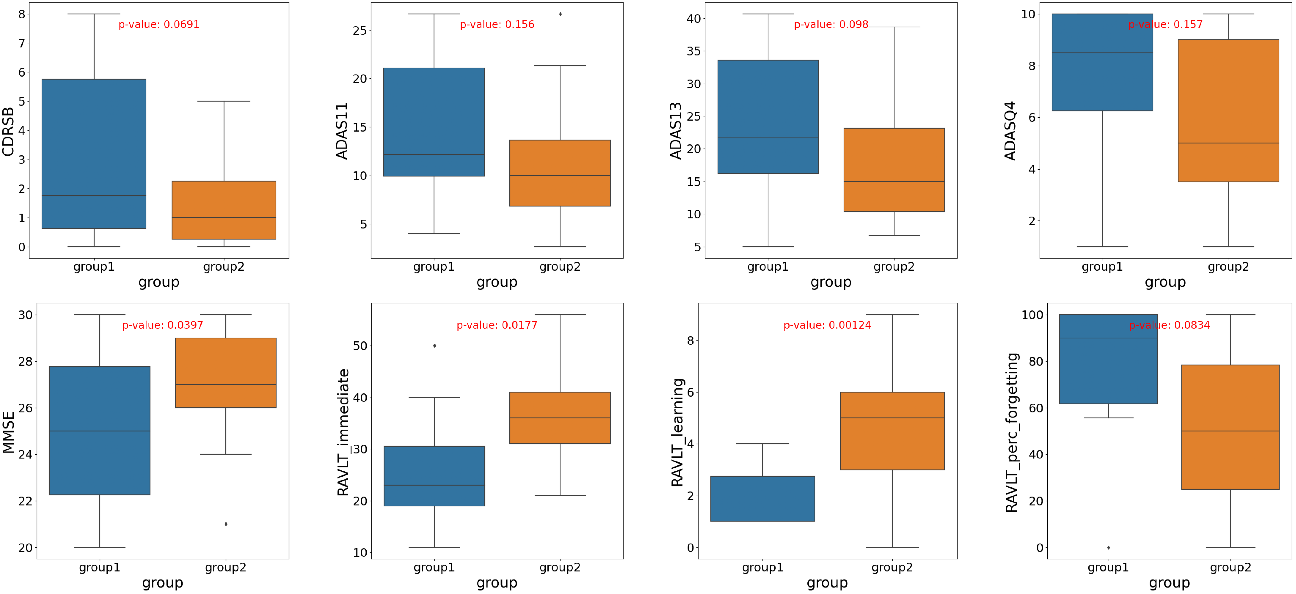
Cognitive Target Score between Group1 and Group2. For every test score, it shows that the cognitive level of Group 1 is lower than Group 2.

## Discussion

In this study, we present a novel model called Het-PR, designed to perform Personalized Regression through the implementation of a Linear Mixed Model. Our proposed model demonstrates superior accuracy in recovering cohorts, finding the personalized genetic basis compared to the current state-of-the-art methods, as validated through an extensive simulation study and real data experiments. In the context of Alzheimer’s disease research, the Het-PR model’s ability to select SNPs proves to be highly reliable, based on two key factors: (1) by verifying whether the selected SNP has previously been identified to be implicated with AD, and (2) more interestingly, showing a first attempt to dividing patients genetic basis based on cognitive levels. The findings of this study suggest that the Het-PR model has the potential to be further employed in determining the stage of Alzheimer’s disease, offering a chance to the field of personalized medicine and Alzheimer’s disease stage assessment.

Although we only conducted the experiment on Alzheimer’s SNP data, it’s worth noting that our model can be applied to any heterogeneous medical data for further research, such as pan-cancer analysis, Asthma’s pathological analysis, depression analysis, etc. The Het-PR model is also potential to be applied in financial, social and other fields.

In the context of this research, several limitations must be acknowledged. First of all, the constrained size of the dataset impedes a definitive understanding of the relationship between cognitive levels and their genetic determinants. As a logical progression, it is imperative to apply the proposed methodology to an expansive Alzheimer’s database to garner more comprehensive insights. Additionally, the current framework, especially concerning the expansion to *σ*^2^ ⊙ *diag*(**XX**^**T**^) as delineated in Equation 14, represents a foundational model. This prompts a series of academic inquiries: How might one adapt *σ* to accommodate individual covariates? Is there a theoretical justification for incorporating the complete matrix? These considerations, among others, necessitate rigorous exploration in subsequent research endeavors.

## Supporting information

**S1 Appendix. Supplementary figures and tables**. The appendix includes the Top 10 influential SNPs figure, etc.

## Acknowledgement

The authors would like to thank Caleb N. Ellington from Carnegie Mellon University for his contribution in parts of the codes and early-stage discussions.

